## Supporting Information for "Kinetic and data-driven modeling of pancreatic β-cell central carbon metabolism and insulin secretion"

**Fig S1. Collected Metabolomics Data.** The fold change direction (increase- blue, decrease-red, or no change-yellow) in 13 published mass spectrometry datasets (*x-axis)* for all metabolites present in the computational model.

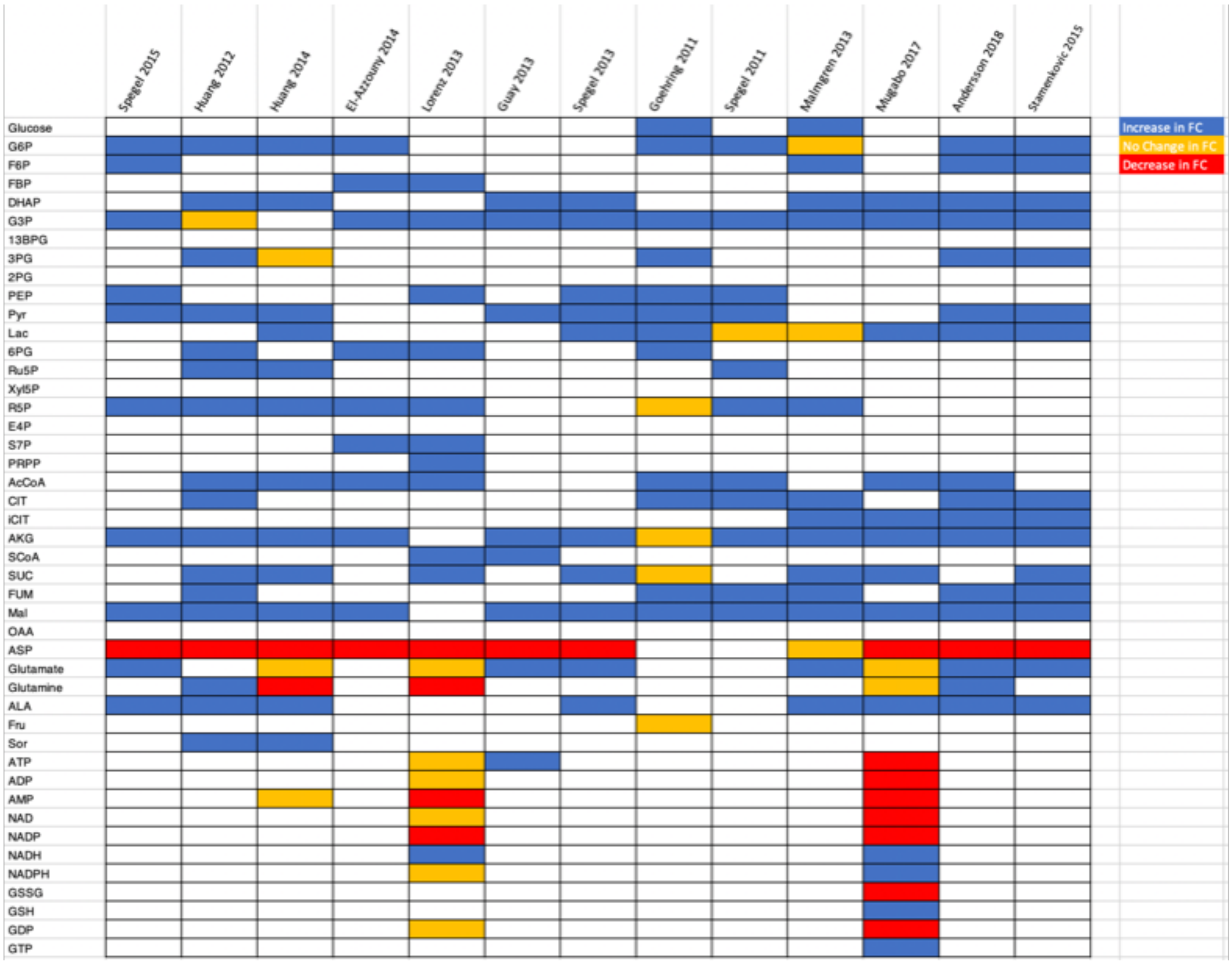

**Fig S2. Estimated Parameter Values.** Having run an eFAST sensitivity analysis, 32 model parameters were found to be influential and were subsequently fit using PSO. The 8 best parameter fits were used for all subsequent analyses and for building the partial least squares regression model. The distribution of fitted model parameter values is shown here.

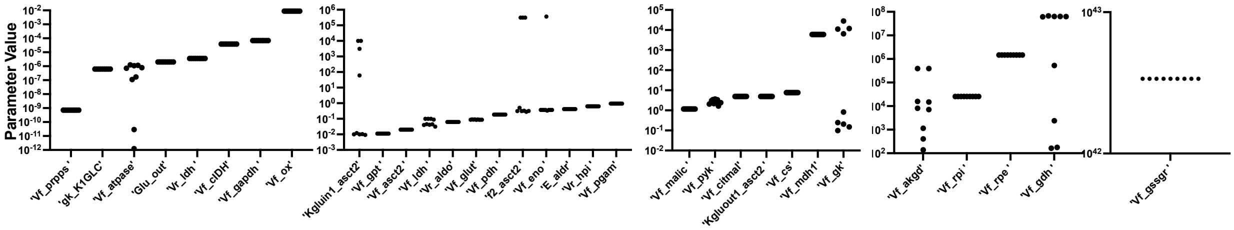

**Figure S3: Effect of *ak* reaction perturbation.** We perturbed the adenylate kinase reaction by increasing its $V_{max}$ value by a factor of 5, and assessed the effect on the network, comparing metabolite levels, reaction fluxes, and insulin secretion to the unperturbed condition.

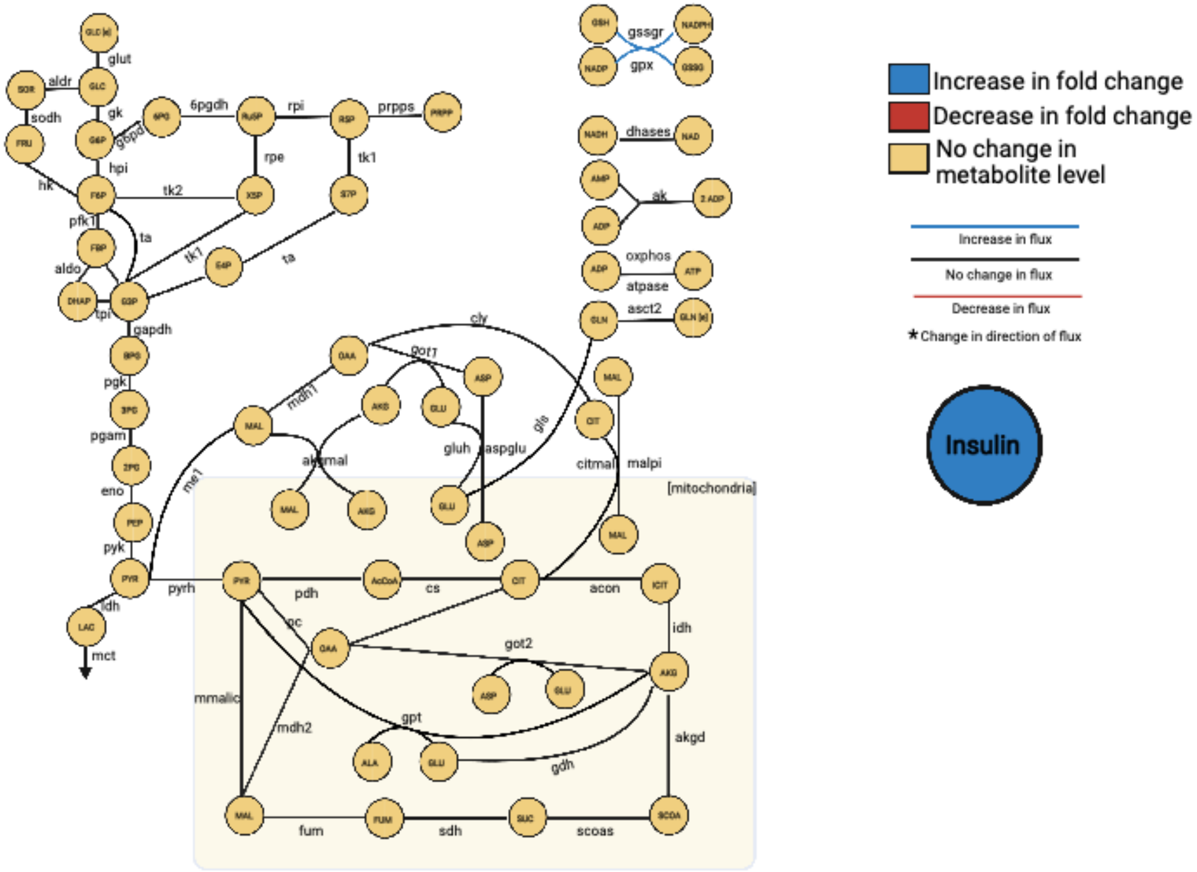
